# Real-time detection of circulating tumor cells in living animals using functionalized large gold nanorods

**DOI:** 10.1101/498188

**Authors:** Rebecca Dutta, Orly Liba, Elliott D SoRelle, Yonatan Winetraub, Vishnu C. Ramani, Stefanie S. Jeffrey, George W. Sledge, Adam de la Zerda

**Affiliations:** Department of Structural Biology, Stanford University, Stanford, CA 94305.; Molecular Imaging Program at Stanford and the Bio-X Program, Stanford, CA 94305.; Electrical Engineering, Stanford University, Stanford, CA 94305.; Biophysics Program at Stanford, Stanford, CA 94305.; Department of Surgery, Stanford University School of Medicine, Stanford, CA, 94305.; Department of Medicine, Stanford University Medicine, Stanford, California; The Chan Zuckerberg Biohub, San Francisco, California 94158, USA

## Abstract

Optical coherence tomography (OCT) with significant speckle reduction can be used with highly-scattering contrast agents for noninvasive, contrast-enhanced imaging of living tissue at the cellular scale. The advantages of reduced speckle noise and improved targeted contrast can be harnessed to track objects as small as 2 μm *in vivo*, with the potential for cell tracking and counting in living subjects. Here we demonstrate the use of Large Gold Nanorods (LGNRs) as contrast agents for detecting individual micron-sized polystyrene beads and single myeloma cells in blood circulation using speckle modulating-OCT (SM-OCT). This is the first time that OCT has been used to image at the individual cell scale *in vivo*. This technical capability presents an exciting opportunity for the dynamic detection and quantification of tumor cells circulating in living subjects.

## Text

The ability to count and track single cells flowing in the bloodstream of a live animal is a central goal in the growing field of circulating tumor cell (CTC) research.^1,2^ CTCs, which are cells that have shed from a primary tumor and entered into the vasculature, may act as seeds for future metastases; thus the enumeration of CTCs in cancer patients has important prognostic and therapeutic value. Most current *in vitro* attempts at enumerating and isolating CTCs such as CellSearch have focused on blood biopsies.^3–5^ However, the rarity of CTCs (~1-10 CTCs/mL of blood), the limit on blood sample volume, and typical sampling frequency virtually precludes the acquisition of dynamic information about the number of tumor cells in the bloodstream using these methods.^6,7^

*In vivo* imaging could allow for dynamic tracking of tumor cells as a diagnostic test as well as a therapeutic test to determine the effect of different drugs on the number of tumor cells in the bloodstream. Over the past few years, the use of contrast agents has advanced the field of *in vivo* optical imaging techniques such as MRI, photoacoustic imaging, and positron emission tomography. However, due to the poor spatial resolution of these techniques, tissue cannot be examined at the single-cell scale.^8,9^ Single-impulse panoramic photoacoustic computed tomography (SIP-PACT) has been used to image CTC clusters in the microvasculature of living mice; however, SIP-PACT does not have the resolution to image at the individual cell level and relies on the intrinsic scattering of specific cell types, such as melanoma cells.^10^ *In vivo* flow cytometry has also been used to detect flowing cells but relies on the use of fluorescent proteins.^11^ Optical coherence tomography (OCT) is a noninvasive, label-free, optical imaging technique that uses low-coherence interferometry to visualize tissue at millimeters depth of optical penetration and subcellular resolution.^12^ OCT is used widely for ophthalmological applications^13^; however, as with all imaging technologies that employ coherence-based detection, speckle noise hinders the diagnostic potential of OCT at the cellular level by significantly reducing spatial resolution.^14^ As our goal is to image small, fast-flowing structures, speckle removal becomes trickier because imaging fast-flowing blood does not enable frame averaging, a technique generally used for speckle removal.^15^ Instead we perform A-scan (or line) averaging which is faster. It allows reduction of time-varying noise, such as photon shot-noise, but does not remove speckle because the imaged volume is static in this small amount of time. Therefore, to reduce speckle, we use SM-OCT. Recently, we have shown that light manipulation through the use of a diffuser can virtually eliminate this speckle noise, enabling OCT to have the unprecedented ability to image individual cells within turbid tissue. Speckle-Modulating OCT (SM-OCT) is able to accomplish this speckle reduction using a ground-glass diffuser that induces local phase shifts in the light illuminating the sample and the light reflected back from scatterers in each imaging voxel.^16^ The result is the removal of speckle noise without degradation of image resolution, allowing for the illumination of structures that were otherwise obscured.

Nanoparticles fine-tuned with respect to different properties such as size and spectral peak can be used to meet specific imaging needs for OCT. Previous work in the lab has taken large gold nanorod contrast agents (LGNRs, ~100 x 30 nm), which backscatter significantly more light than smaller gold nanorods, and biostabilized them so that they can be utilized *in vivo*. LGNRs are coated with poly(sodium 4-styrenesulfonate) (PSS) for stabilization and can then be incubated with living cells for nonspecific uptake of contrast agent; LGNRs can also be further functionalized through the bioconjugation of cell-specific antibodies like anti-epithelial cell adhesion molecule (anti-EpCAM).^17,18^

With the use of improved contrast agents and speckle reduction, the spatial resolution inherent to OCT can be harnessed to visualize contrast agent-labeled objects in flowing blood and at a cellular scale. This new capability is first demonstrated by imaging LGNR-labeled 2 μm polystyrene beads, which have been intravenously injected into a living mouse, circulating in the pinna vasculature. Then, the number of labeled beads detected at a cross section of a blood vessel in the mouse ear are counted temporally to determine their circulation time. Next a similar capability is demonstrated for imaging intravenously injected RPMI-8226 myeloma cells, pre-incubated with LGNRs, as they flow through blood vessels. Figure 1 shows the schematic of the experimental setup for the imaging of injected LGNR-labeled cells in the blood vessel of a living mouse.

**Figure 1.**
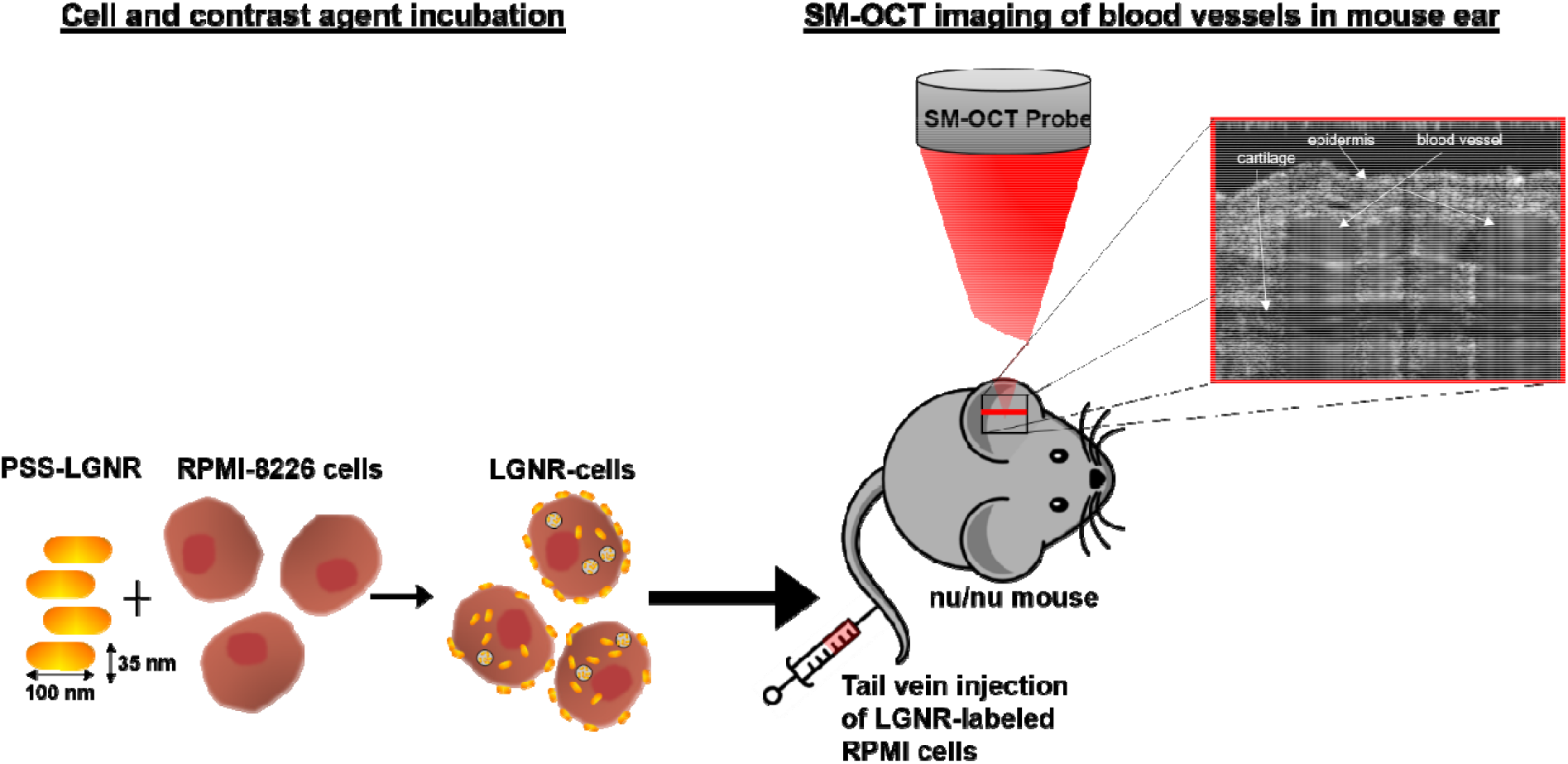
Schematic of experimental process. **A)** LGNRs biofunctionalized with a coating of PSS are incubated with RPMI-8226 cells, resulting in LGNR-labeled cells. **B)** Incremental injections of labeled cells are administered into the tail vein of the mouse. SM-OCT scans at the cross section of blood vessels found in the ear of the mouse, and multiple scans are acquired at the same location to determine the number of cells flowing through the blood vessel over time.

Objects in the bloodstream that take up less volume than a single voxel and exhibit poor scattering need to be adequately labeled with a sufficient number of LGNRs for sufficient enhancement from the LGNRs’ increased scattering and subsequent visualization with SM-OCT. To demonstrate this, 2 μm biotinylated polystyrene beads which are smaller than a single imaging voxel and can be incubated with Neutravidin-functionalized LGNRs are first used for labeling. SEM images indicated that each bead had an average of 159 LGNRs attached non-homogeneously to the surface after excess LGNRs were washed off, surpassing the system’s detection sensitivity (Fig. 2a,b). Unlike unlabeled beads, which are not visible in a PBS suspension with SM-OCT, the labeled beads are individually visible, indicating that objects previously invisible to SM-OCT could be illuminated with adequate LGNR labeling. A suspension of LGNR-beads in PBS was then imaged in capillary tubes, and, unlike the unlabeled beads, individual labeled beads could be visually identified, indicating that objects previously invisible to SM-OCT could be illuminated with adequate LGNR labeling (Fig. 2c). The SM-OCT signal (on linear scale) from contrast agent-labeled beads was two-fold higher than that from unlabeled beads (Fig. 2d). In a separate experiment, labeled and unlabeled beads were imaged in whole mouse blood inside capillary tubes.. Unlabeled beads, which were visible in PBS, were no longer detectable in blood; however, the LGNR-labeled beads were detectable in blood and had a higher SM-OCT signal compared to the surrounding blood. (Fig. 2e). The number of beads detected in a single frame (only considering the top half of each tube, which is less influenced by blood absorption) was representative of the initial concentration of beads that were mixed into the whole blood. As the ear of a mouse is approximately 100 μm thick and the top half of the capillary tube is 200 μm, this did not affect subsequent *in vivo* imaging. The SM-OCT signal of the labeled beads was compared to regions of blood in the blood-only capillary tube that were at the same depth, to account for attenuation of light from red blood cells. When the SM-OCT signal was normalized for depth-dependent attenuation of signal, the signal-to-background ratio (SBR) of labeled beads to blood is 6.3 ± 1.0 (Mean ± SD) fold higher.

**Figure 2.**
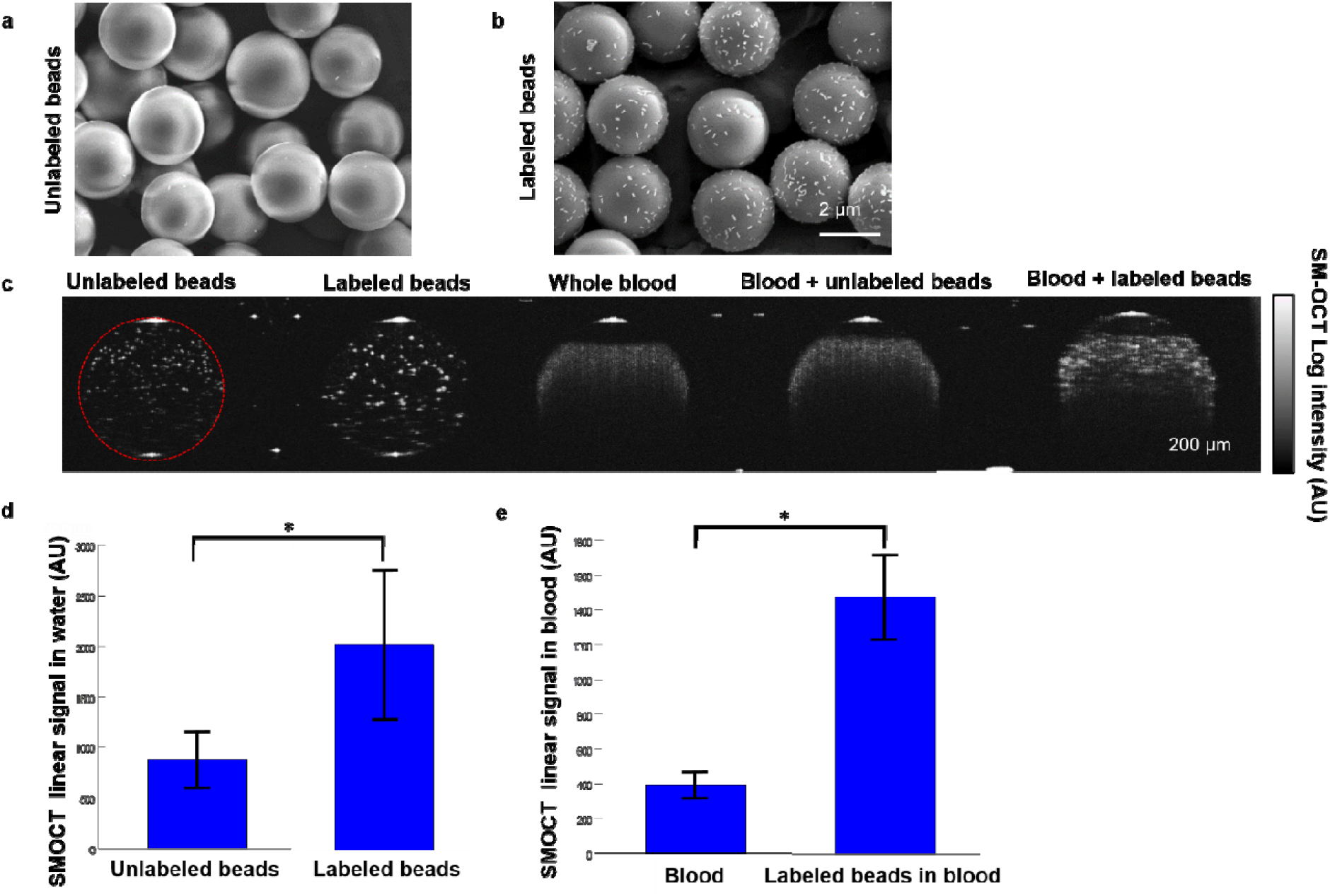
LGNR-labeled beads have a increased signal to background ratio compared to nonlabeled beads. SEM images of A) unlabeled polystyrene beads and B) labeled polystyrene beads. Labled beads had 158.53 ±82.7 (Mean ± SD) LGNRs per bead. C) SM-OCT image of beads in water, labeled beads in water, whole blood, whole blood +unlabeled cells/beads, or with whole blood + labeled cells/beads. Log intensity images were created during post processing. D) Labeled beads have more than 2-fold stronger SM-OCT signal than unlabeled beads (*p<0.001). E) Labeled beads in blood have more than 3-fold more signal than the signal from whole blood without beads (*p<0.001). Unlabeled beads have too weak SM-OCT signal to be spotted in blood and were excluded from this data.

To mimic blood flow, a programmable syringe pump was used to pass whole blood with labeled beads through a capillary tube at biologically relevant speeds. We were able to detect labeled beads in whole blood flowing through the capillary tube at 10 cm/s and 20 cm/s, proving that labeled beads can be detected with SM-OCT at different flow speeds (Fig. 3 a,b). To confirm detection of labeled beads in a background of flowing blood, time-lapse visualization of SM-OCT signal was performed at the location of the punctate signal that was suspected to be a flowing bead. It was found that flowing LGNR-beads had a distinctly different SM-OCT signal pattern over consecutive A-scans than background signal of blood, with background signal showing low, disparate signal and the punctate signal showing a more predictable localized pattern over the time it takes for the bead to leave the imaging region (Fig. 3 c-e).

**Figure 3.**
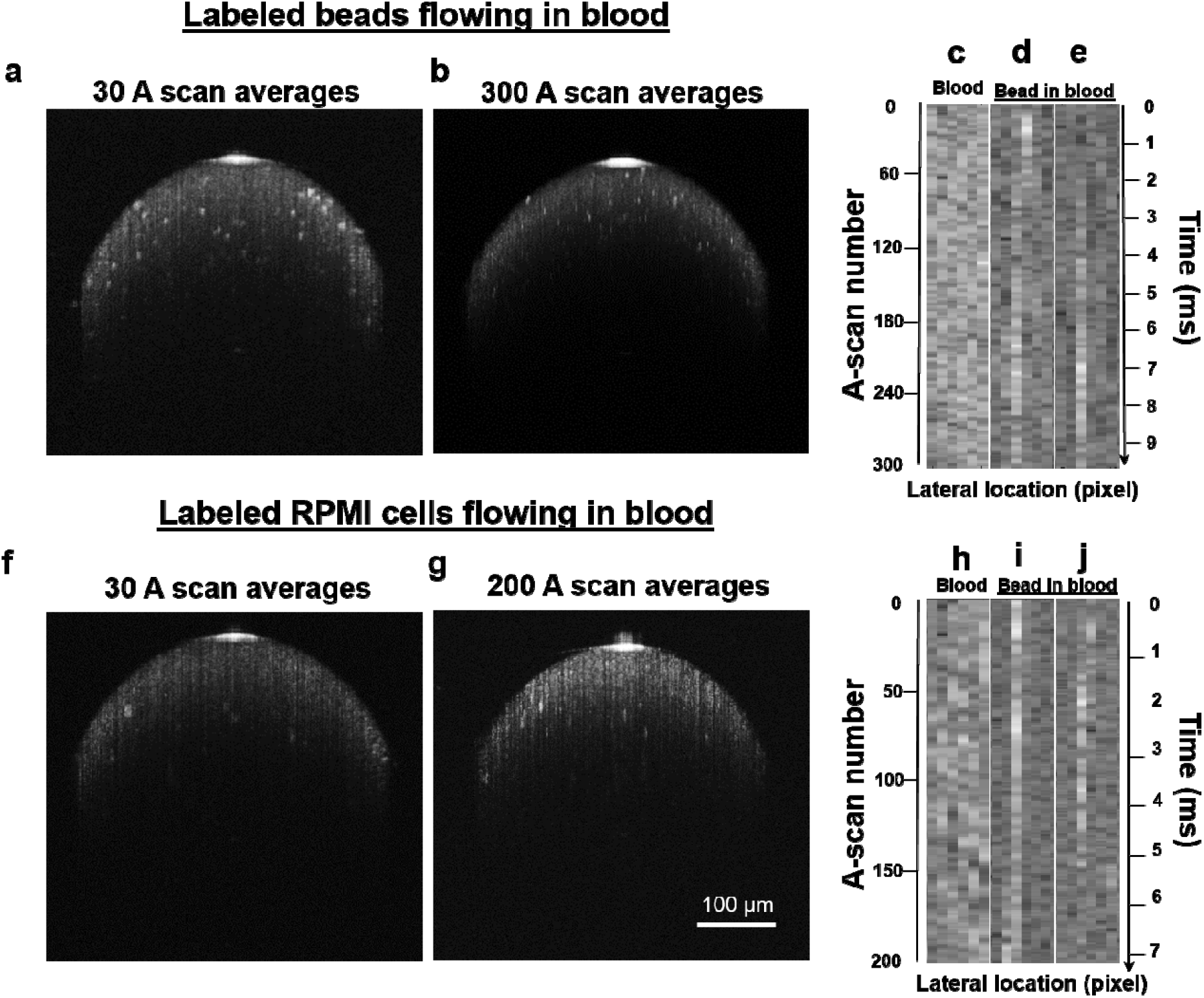
Flowing LGNR-labeled beads and cells have a characteristic SM-OCT signal allowing for clear distinction in blood. Log SM-OCT signal of labeled beads flowing at 20 cm/s through a capillary tube filled with whole blood with A) 30 A-scan averages and B) 300 A-scan averages. A visualization of A-scans was made at adjacent locations through time, showing the linear SM-OCT signal of C) linear SM-OCT signal from flowing whole blood, D,E) beads in the 300 A-scan average imaging session and their immediately adjacent pixels. Frames capturing labeled cells flowing at 20 cm/s through whole blood with F) 30 A-scan averages and G) 200 A-scan averages. Once again, visualization of A-scans was made at adjacent locations through time, showing the linear SM-OCT signal of H) flowing whole blood background, and I,J) 2 cells in the 200 A scan average imaging session and their immediately adjacent pixels.

Beads labeled via the same procedure as in the capillary tube experiments were injected intravenously into the tail vein of a mouse. Image acquisition at the blood vessels of the mouse pinna was done every two minutes post-injection for 30 minutes, at which point, no beads could be detected in the blood vessel. Labeled beads showed up in the blood vessel as high-intensity puncta (Fig. 4a). The puncta pattern changed with every frame since the blood flow within the blood vessel is faster than the image acquisition rate. These punctate signals were similar in size and signal intensity to those seen in the capillary tube imaging experiments. To estimate the circulation time of the beads, the punctate signals inside the vessels were manually counted in each frame. The number of puncta was graphed over time (Fig. 4b). Sixteen minutes post-injection, the number of puncta counted over 100 consecutive frames was found to be approximately similar to the number of puncta counted pre-injection, indicating that the labeled beads had largely been cleared from the circulation (two-tailed t-test, p < 0.05).

**Figure 4.**
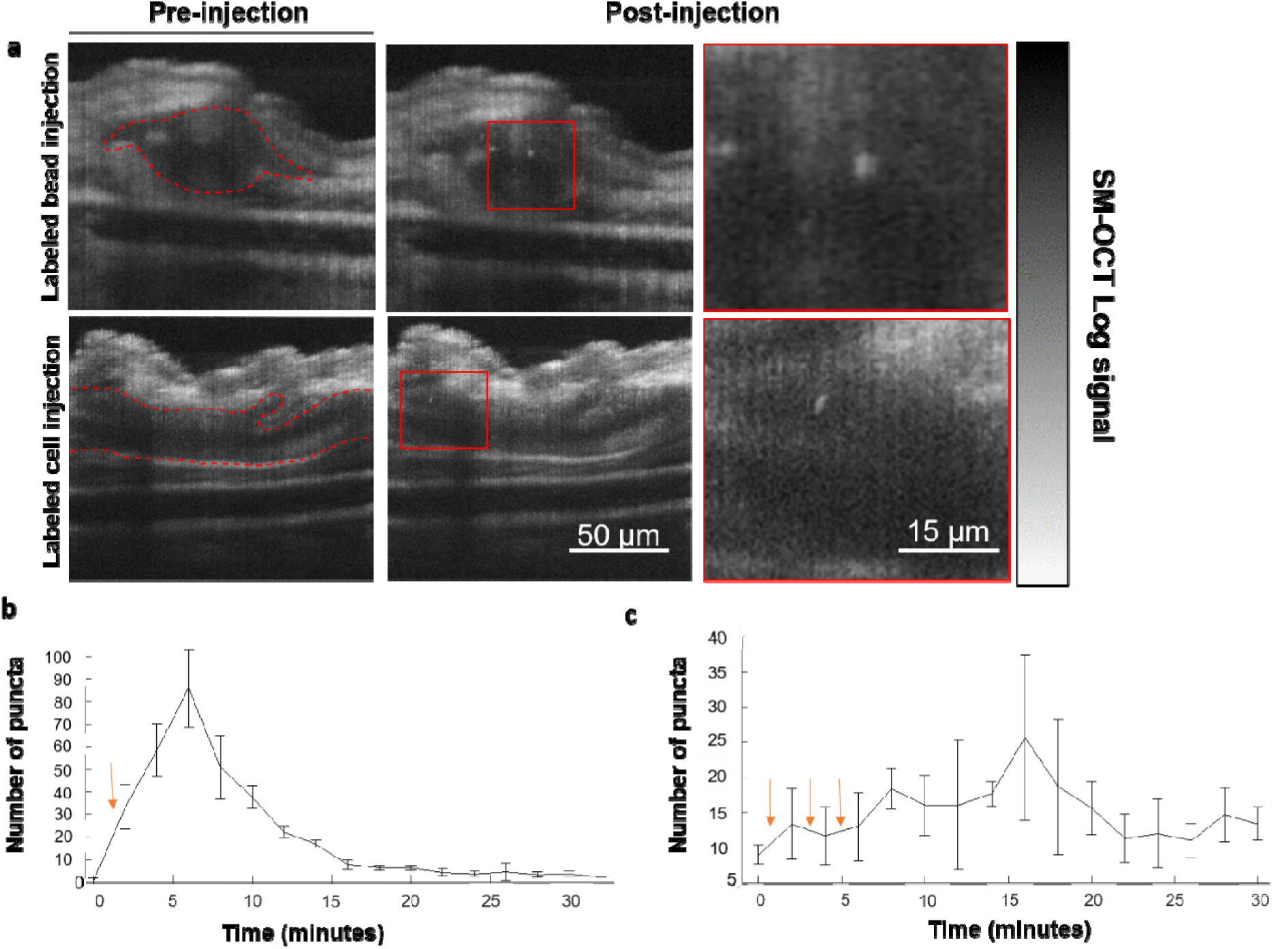
Labeled beads and labeled RPMI-8226 cells are detected in the bloodstream post intravenous injection. A) Top panel: strong punctate signal is seen in the cross-section of a blood vessel of the ear pinna post-injection, indicating a passing labeled polystyrene bead. Red dotted line indicates boundary of the blood vessel of interest. Bottom panel: longitudinal cross section of a blood vessel pre and post-injection reveals the presence of an increased number of punctate signal similar to the one indicated in the red box that can be attributed to flowing cells labeled with LGNRs. B) Number of punctate signals in the cross section of the blood vessel were counted over a 30 minute period, at 2 minute intervals, to track circulation time of labeled beads (mean ±SD). Red arrow indicates time of injection. C) Number of punctate signals in the cross section of the blood vessel were counted over a 30 minute period, at 2 minute intervals, to track circulation time of labeled cells (mean ±SD). Red arrows indicate times of incremental injections.

Given the ability to detect circulating sub-cellular-sized labeled polystyrene beads using SM-OCT, we proceeded with labeling cells derived from tissue culture with LGNRs to determine their detection in blood vessels of living animals. Due the difficulty of obtaining and propagating primary cultures of CTCs from human patients^19–21^, we used cells from the RPMI-8226 myeloma cell-line, which is grown as a suspension culture and well-suited with regard to its circulation mechanics, to tumor cells in circulation, as suggested by its similarity to circulating white blood cells.^22^ Additionally, unlike cancer cells of epithelial origin, myeloma cells do not have a lot of cell surface integrins and therefore have low cell-cell or cell-ECM adhesion^23^ that would encourage adherence to blood vessel walls. RPMI-8226 cells were incubated for one hour with an excess of LGNRs and washed thoroughly to remove any unbound LGNRs. To verify LGNR labeling of RPMI-8226 cells, labeled cells and unlabeled cells was imaged using a hyperspectral dark-field microscope with a range of 400-1000 nm. A custom machine-learning algorithm for adaptive detection of LGNRs based on their unique optical signature^24^ was used to identify the location of the LGNRs in the RPMI-8226 cells. The majority of cells incubated with LGNRs show a significant amount of pixels that have strong scattering in the near-infrared region, suggesting the presence of LGNRs, particularly on the edges of the cell. Pixels that were classified as containing LGNRs are shown in orange on the hyperspectral image (Fig. 5a). As expected, the control sample of unlabeled cells show no presence of this LGNR scattering spectrum.

**Figure 5.**
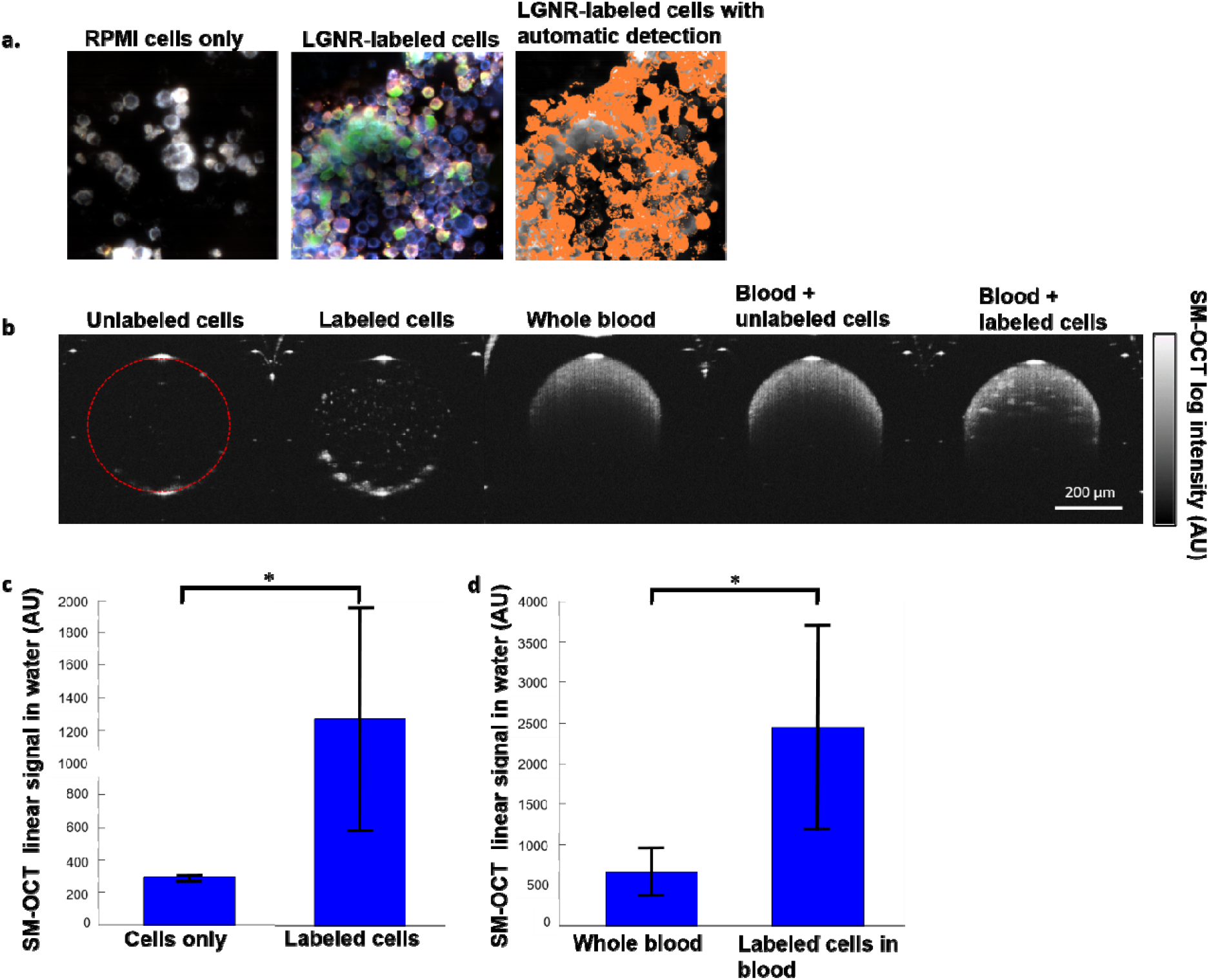
RPMI-8226 cells can be visualized in PBS and whole blood after incubation with PSS-LGNRs. A) Hyperspectral images of the LGNR-labeled and unlabeled cells. Custom algorithms allow for the automatic detection of LGNRs taken up by the cells. Detected locations of LGNRs are shown in orange. B) 2μm polystyrene beads were incubated with LGNRs, washed, and then spiked into whole blood. Capillary tubes were filled with unlabeled cells in PBS, labeled cells in PBS, whole blood, whole blood + unlabeled cells, or with whole blood + labeled cells. Log intensity images were created during post processing. C) Labeled cells have more than 3 fold higher signal than cells alone (p<0.001) and D) more than 2 fold higher signal than whole blood (p<0.001).

A suspension of labeled cells and unlabeled cells in complete cell culture media were imaged in adjacent capillary tubes (Fig. 5b). While single unlabeled cells generally had no detectable SM-OCT signals due to poor scattering of light compared to their surroundings, cells labeled with LGNRs could be clearly distinguished in the capillary tube owing to the enhanced scattering from the LGNR contrast agents. LGNR-labeled RPMI-8226 cells had three-fold higher SM-OCT signal compared to unlabeled cells (Fig. 5c).

Labeled cells were then added into whole mouse blood to determine whether the signal intensity of the labeled cells was higher than the baseline intensity from the whole blood. Unlike unlabeled cells, LGNR-labeled RPMI-8226 cells in whole blood could be visualized in top half of the capillary tube (bottom 200 μm were not analyzed because of attenuation from blood)

(Fig.5b). The signal from labeled cells was compared to the signal from blood at similar depths, and it was found that the LGNR-labeled RPMI-8226 cells had two-fold greater signal than blood (Fig. 5d). Moreover, similarly to the labeled beads, the signal from labeled cells was highly punctate, allowing for their clear demarcation in a background of lower backscattering. These results indicated that RPMI-8226 cells can be sufficiently labeled with LGNRs such that their light backscattering increased enough to allow for their detection by SM-OCT in blood vessels.

To simulate circulation of labeled cells in blood circulation, LGNR-labeled RPMI-8226 cells in whole blood were passed through a capillary tube using the automatic syringe pump. Similar to the labeled beads described earlier, the labeled cells were detectable flowing at 10 cm/s as well as 20 cm/s, indicating that labeled cells can be detected with SM-OCT in a range of biologically relevant flow speeds (Fig. 3 f,g). To confirm detection of labeled cells in a background of flowing blood, a time-lapse analysis was performed of the SM-OCT signal at the location of the punctate signals that were suspected to be flowing cells (Fig. 4 h-j). As with the beads, flowing LGNR-cells had a distinctly different SM-OCT signal pattern over consecutive A-scans compared to the background signal. Importantly, the number of LGNR-cells counted in any one frame corresponded to the concentration of cells that were added into whole blood. We predicted a count of 37.8 beads in every frame, and found that we could count 17.4± 5.4 (Mean± SD), or approximately half of the expected bead count. This count makes sense because only the top half of the capillary tube can be analyzed due to the attenuation of light by the red blood cells and indicates that the scan speed of SM-OCT is fast enough to capture every cell flowing through the cross-section of the capillary tube at flow speeds up to 20 cm/s.

Next, we tested our detection scheme in the bloodstream of a live mouse. RPMI-8226 cells were prepared and incubated with LGNRs using the same method as in the capillary tube experiments. Approximately five million cells were injected incrementally to the mouse’s tail vein to allow for sufficient clearance of cells through the capillary vessels of the lungs. Imaging was performed after each injection increment at the longitudinal cross-section of a blood vessel in the pinna of the mouse. Lengthwise sections were imaged at the middle of the blood vessel since the probability of having a cell in an individual cross-section is rare and a lengthwise slice allows for the sampling of a much larger region of the blood vessel at any given time. Bright-field microscopy of blood collected via cardiac puncture from a mouse injected with labeled cells showed that less than 1% of the 1095 cells counted are not single cells. Of the 10 multi-cellular clumps that were counted in the sample, 9 were clumps consisting of 2 RPMI-8226 cells, and 1 was a clump of 3 RPMI-8226 cells. If all five million of the cells injected made it into circulation, punctate signals should be seen around once every 8 frames in a blood vessel that is approximately 50 μm in size; however, bright punctate signals were spotted in the blood vessel approximately once every 14 frames, suggesting that less than half of the cells injected make it into systemic circulation. If all 1% of cells that are multicellular clumps make it into systemic circulation, 97% of cells detected would still be single cells. This strongly suggests that the cells in circulation that are detected by SF-OCT are, indeed, single cells and not clusters of cells.

Bright punctate signal considered to be LGNR-labeled RPMI-8226 cells were manually counted every two minutes over a 30 minute period. As with the beads, there was an increase in punctate regions after the injection, indicating the presence of LGNR-labeled RPMI-8226 cells flowing *in vivo* (Fig. 4a). The ability to detect these cells in the blood vessel of a living mouse once they have been sufficiently labeled with the LGNR contrast agents demonstrates that SM-OCT has a sufficient resolution to be able to resolve circulating cells. The number of punctate signals counted were graphed over time to better understand the circulation dynamics of the injected cells (Fig. 3c). Unlike the rapid circulation and clearance of labeled beads in the bloodstream, labeled cells showed a less predictable circulation pattern, with a slower, more gradual increase in the number of punctate signals counted over time as well as a more gradual return to baseline. There are several possible explanations for this. While beads do not have a tendency to stick to vasculature, cells needed to be injected incrementally to prevent a backup of cells in pulmonary microcirculation which could potentially lead to the death of the mouse. Prior research has noted the limited clearance of intravenously injected tumor cells that arrest in the pulmonary vasculature^25^. Coupled with the distal location of the ear pinna blood vessel, the circulation pattern of cells becomes more stochastic and depends on a variety of factors such as the tendency of the cells to adhere to vessels and their deformability.

During analysis of pre-injection frames, rare endogenous features with high signal intensity punctate, similar to that of flowing LGNR-labeled beads or labeled cells, were noticed. The presence of punctate signal due to possible scatterers present in the blood or on the surface of the blood vessel must be distinguished from the flowing labeled beads or cells of interest to reduce the number of false positive signals counted. During cell/bead counting, we strictly avoided any punctate signals that occurred repeatedly in the same location in both pre and post-injection images. It is likely that these signals are the result of backscattered light from highly scattering cells on the surface of the blood vessel; flowing cells, given their rarity, are less likely to show up in the same place in several frames. Further signal analysis, including possible characterization of the spectral characteristics of these punctate signals, could better distinguish between cells of interest and highly-scattering species that are endogenous to the mouse.

Using highly scattering LGNRs to enhance the spectral signal of target objects allows us to use SM-OCT to identify individual sub-cellular sized beads as well as tumor cells as they flow through the blood vessel of a living mouse. Unlike conventional spectral-domain OCT, which would not allow for accurate single-cell detection due to high amounts of speckle, the SM-OCT setup allows us to distinguish moving cells from speckle noise. In addition, the use of LGNRs, which have higher backscattering compared to smaller GNRs, allows us to distinguish flowing labeled cells and beads from the background signal of blood.

Here we have demonstrated the ability to track objects as small as 2 μm within the circulation of a living mouse. This result has the potential for impacting research in cell tracking, particularly in the tracking of tumor cells in circulation *in vivo* and enable the research of CTC extravasation, migration, and seeding. In this report, tumor cells are seen clearly flowing in the bloodstream noninvasively using SM-OCT. This effort is maximized by the use of highly scattering LGNRs as a contrast enhancer for low-scattering tumor cells as well as the removal of speckle noise in OCT to increase effective resolution. With additional biofunctionalization of LGNRs, intravenously injected contrast agents could be targeted to CTC- or cancer cell-specific cell surface antigens such as EpCAM or human epidermal growth factor receptor 2 (HER2), enabling targeted attachment to possible metastatic cells. Using SM-OCT, a patient’s blood could be monitored over time for strong punctate signals, tracking CTCs in the bloodstream and opening up the possibility of more robust treatment plans. The concepts shown here can be extended to dynamically monitor CTCs in patients and may have the potential to be used in addition or as a replacement to blood biopsies.

## Methods

### Labeling Polystyrene beads and preparation for injection

Neutravidin-functionalized polystyrene microbeads with a nominal size of 2.0-2.4 μm (0.5% w/v, Spherotech, Inc., CAS# NVP-20-5) were incubated at room temperature for 90 minutes with an excess of Biotinylated Large Gold Nanorods (LGNRs) with peak absorption at 835 nm and size of approximately 100 x 30 nm. A majority of the excess rods were washed off by centrifuging the coated beads thrice at 900 x g for 1 minute. For programmed syringe flow experiments, 13*10^6 polystyrene beads were incubated with 10 uL of 2.5nM biotinylated LGNRs for 90 minutes and then washed. At the end of the washing, labeled beads were counted using the hemocytometer, and the 5.02*10^5 beads that still remained post-wash were pelleted and diluted in 460 μL of whole blood. For *in vivo* imaging studies, an injection volume of 230 μL at 10^6^ LGNR-labeled beads/μL in PBS was prepared.

### LGNR functionalization

Large Gold Nanorods (LGNRs) were produced using a method described previously.^10^ LGNRs at 835 nm surface plasmon resonance were used because their scattering to absorption ratio is higher than that of smaller gold nanorods. LGNRs were incubated with 100 M poly(sodium 4-styrenesulfonate) (PSS, MW ~ 70 kDa, Aldrich, CAS# 25704-18-1) for 5 minutes with vortexing to produce PSS-LGNRs. LGNRs-PSS were then centrifuged at 2550 x g and resuspended. PSS coating was repeated three times. For bead incubation experiments, PSS-LGNRs at 1nM were incubated with Biotin-PEG-SH (MW ~5 kDa, NANOCS, PG2-BNTH-5k) to produce Biotin-PEG-PSS-LGNRs. Before incubation with either cells or beads, LGNRs were spun down and resuspended to 2.5 nM.

### RPMI-8226 Cell culture and cellular uptake of LGNRs

RPMI-8226 myeloma cell line (Sigma-Aldrich) was cultured under standard conditions using RPMI-1640 medium at 10% fetal bovine serum. For labeling experiments, cells were resuspended in fresh, RPMI-1640 media and then incubated at 36°C with an excess of PSS-coated LGNRs (~2000 LGNRs/cell). After 50 minutes of incubation, cells were washed three times with PBS to remove excess LGNRs. Before being used, labeled cells were vortexed gently for 10 seconds and then passed through a 31 G insulin syringe twice to break up any clumps. Cells were then imaged immediately or prepared for intravenous mice injections. For capillary tube imaging studies, 250 μL of LGNR-labeled RPMI-8226 cells were prepared for injection at a concentration of 20,000 cells/μL in PBS. Cells were kept at 36°C until the start of the injection to ensure viability and retention of LGNRs.

### *In vivo* imaging experiments

Female nude mice (nu-/nu- 6-8 weeks old, Charles River Labs, Wilmington, MA) were used for all imaging studies. Mice were anesthetized at 2% isoflurane and placed on a 37°C heating pad for the duration of the experiment. The mouse ear pinna was mounted on a cuvette using double sided tape to flatten the ear surface. A thin layer of ultrasound gel was applied to the top of the ear, and a cover glass was placed over the gel to optimize imaging. Pre-injection images were taken at the cross section of major blood vessels (50-75 μm) of the ear. Tail vein injections were used to administer either the LGNR labeled polystyrene beads or the LGNR labeled RPMI-8226 cells. For labeled bead injections, the entire injection volume was injected at once and allowed to circulate for 1 minute before images were taken. SM-OCT images were acquired in one minute intervals for the first 5 minutes and then at 5 minute intervals until beads cleared out at approximately the 30 minute mark. 30 A-scan averages were taken over the course of 100 frames for every time point. This ensured that beads were not counted twice and that beads were not missed because the scan rate was slower than the flow velocity through the scan area. Labeled cell injections were made in 50 μL increments to prevent cells from arresting in the lung capillary vessels. From one of the mice injected with LGNR-labeled RPMI-8226 cells, blood was collected using cardiac puncture and then analyzed under confocal microscopy for the presence of RPMI-8226 cells. Blood was kept in heparinized tubes until imaging was performed. The mouse on whom the cardiac puncture was performed was euthanized 3 minutes after the completion of the cell injection to allow for sufficient circulation of injected cells. All animal experiments were performed in compliance with IACUC guidelines and with the Stanford University Animal Studies Committee’s Guidelines for the Care and Use of Research Animals. Experimental protocols were approved by Stanford University’s Animal Studies Committee (APLAC protocol 27499, 28117).

### SM-OCT setup and explanation

**Experimental setup** SM-OCT was implemented by modifying an existing OCT systems: the Ganymede HR (Thorlabs), which is a spectral domain OCT system. The light source of the Ganymede HR is a super luminescent diode (SLD) with a center wavelength of 900 nm. The spectrometer has a 200 nm bandwidth (*l*=800-1000 nm), which provides 2.1 mm axial resolution in water. The spectrometer acquires 2048 samples for each A-scan at a measured rate of 20.7 kHz. All image reconstruction and analysis was performed with Matlab (Mathworks) using raw data from the spectrometer. The first lens of the imaging system (LSM03-BB, Thorlabs) provides a lateral resolution of 8 mm (full-width at half-maximum, FWHM) and depth of field (DOF) of 143 mm in water. A diffuser (DG10-1500-B) was placed at the original focal plane of the OCT probe, and a new focal plane was projected by a 4f imaging system. The 4f configuration was implemented using two similar lenses (LSM02-BB, Thorlabs) that provide a lateral resolution of 4.2 mm (FWHM) and DOF of 32 mm in water. Due to the extension of the sample arm and the addition of 2 lenses and the diffuser, the reference arm was extended by approximately 10 cm, and dispersion compensation elements were added (two LSM02DC, Thorlabs). The reference arm was extended by placing metal rods between the OCT probe and the reference mirror. Conventional OCT images were obtained with the SM-OCT apparatus without the diffuser. In this configuration, light propagates through the 4f imaging system and the extended reference arm. OCT images obtained this way are of similar quality to the OCT images obtained by the original OCT probe, without any modifications. The diffuser was placed in the focal plane of the first lens and held by a mount within a rotating motor (RSC-103, Pacific Laser Equipment). To achieve maximal diffuser speed at the location of the beam, it was focused near the outer edge of the diffuser, where the velocity was approximately 9 mm s^-1^. This velocity was sufficient to create decorrelated speckle patterns in each A-scan^16^, therefore, in this setup, the number of uncorrelated speckle patterns is equal to the number of acquired A-scans.

## Acknowledgements

This work was funded in part by grants from the Susan G. Komen Breast Cancer Foundation (SAB15-00003), Claire Giannini Fund United States Air Force (FA9550-15-10007), the National Institute of Health (NIH DP50D012179), the National Science Foundation (NSF 1438340), the Damon Runyon Cancer Research Center (DFS# 06-13), the Mary Kay Foundation (017-14), the Donald E. and Delia B. Baxter Foundation, the Skippy Frank Foundation, a seed grant from the Center for Cancer Nanotechnology Excellence and Translation (CCNE-T; NIH-NCI U54CA151459), and a Stanford Bio-X Interdisciplinary Initiative Program (IIP6-43). A.d.l.Z is a Chan Zuckerberg Biohub investigator and a Pew-Stewart Scholar for Cancer Research supported by The Pew Charitable Trusts and The Alexander and Margaret Stewart Trust. O.L. is grateful for a Stanford Bowes Bio-X Graduate Fellowship. E.D.S. wishes to acknowledge funding from the Stanford Biophysics Program training grant (T32 GM-08294). We wish to thank Peng Si for his help with the SEM, Rakhi Gupta for cell culture support, and Elias Godoy with his help obtaining mouse blood for imaging experiments.

